# A neurodevelopmental origin of behavioral individuality

**DOI:** 10.1101/540880

**Authors:** Gerit Arne Linneweber, Maheva Andriatsilavo, Suchetana Dutta, Liz Hellbruegge, Guangda Liu, Radoslaw K. Ejsmont, Lisa M. Fenk, Andrew D. Straw, Mathias Wernet, Peter Robin Hiesinger, Bassem A. Hassan

## Abstract

The genome *versus* experience, or “Nature *versus* Nurture”, debate has dominated our understanding of individual behavioral variation. A third factor, namely variation in complex behavior potentially due to non-heritable “developmental noise” in brain development, has been largely ignored. Using the *Drosophila* vinegar fly we demonstrate a causal link between variation in brain wiring due to developmental noise, and behavioral individuality. A population of visual system neurons called DCNs shows non-heritable, inter-individual variation in right/left wiring asymmetry, and control object orientation in freely walking flies. We show that DCN wiring asymmetry predicts an individual’s object responses: the greater the asymmetry, the better the individual orients. Silencing DCNs abolishes correlations between anatomy and behavior, while inducing visual asymmetry via monocular deprivation “rescues” object orientation in DCN-symmetric individuals.

**One Sentence Summary:** Non-heritable individual variation in neural circuit development underlies individual variability in behavior.

## Main Text

Individual variability in external and internal organ morphology is highly abundant in all organisms, including among genetically identical individuals, such as human identical twins and species that reproduce by parthenogenesis (*1-3*). In this regard, the brain is no exception. The simplest examples of individual brain variation include differences of size and weight of human brains (*4*) but also variance of more complex traits, like neuroanatomical parcellations, have been described (*5, 6*). The same variability is also found at a deeper organizational level in the number of neurons (*7*). In invertebrate model organisms, it was shown that individual neurons show varying morphology and wiring (*8*), variable and plastic synaptic morphology and molecular composition (*9-12*).

The same level of individual variability can also be found in behavior, the main output of nervous system function (*13*). Complex innate behaviors, like selective attention to stimuli, show significant individual variation even amongst genetically similar or even identical individuals (*14-16*). Stability of these individual differences over time allows to define these behavioral idiosyncrasies as animal individuality (*17, 18*). Analysis of neural circuits in various genetic model organisms has led to the widely-held view that variability in innate behavior is largely due to neuromodulation of otherwise anatomically hard-wired neuronal circuits (*19, 20*). This includes stable behavioral traits in *C. elegans* foraging that depend on serotonin modulation (*19*) and serotonergic and dopaminergic regulation of the zebrafish larval startle response (*20*). More recently it was shown that the developmental plasticity of higher order neural circuits, partly driven by stochastic mechanisms, makes them intrinsically variable, resulting in a range of possible circuit diagrams even amongst genetically identical individuals. This variability in developmental wiring is regulated by cell-cell signaling events and results in differential gene expression profiles amongst otherwise indistinguishable neurons (*21, 22*). However, despite much interest, debate and speculation, the contribution of non-heritable developmental variability in neural circuit wiring to individual behavioral variation remains almost completely unexplored (*23, 24*).

To test whether probabilistic wiring of neural circuits affects behavioral variation, we used a *Drosophila* higher order visual circuit, called the Dorsal Cluster Neurons (DCN) (*25*). DCNs exhibit up to 30% wiring variability of their axonal projection not only between individuals, but also between the left and right hemispheres of the same brain (*26*). DCNs in each hemisphere derive from a single neural stem cell (*25, 27*) and their axons innervate two alternative target areas in the fly visual system called the lobula and the medulla, respectively (*25*). The medulla processes visual motion information (*28*), while the lobula is implicated in the integration of visual and motor information. Initial characterization of DCN anatomy showed that about 25% of DCN axons have their terminal presynaptic arbors in the contralateral medulla (M-DCNs), while about 75% have their terminal presynaptic arbors in the contralateral lobula (L-DCNs). L-DCN axons also project onto M-DCN dendrites, while M-DCN axons project onto medulla neurons. Because the decision that each DCN makes between being M-DCN or L-DCN is determined by an intrinsically stochastic lateral inhibition mechanism (*21*).

In order to test a quantitative relationship between a behavior and non-heritable brain wiring variability, a visual behavioral assay was required that allows testing individual flies repeatedly over a significant stretch of time. For this purpose, the robust and easily accessible Buridan’s paradigm was chosen (*29*). In this assay a single fly walks between two equally attractive visual targets in the form of high contrast black stripes placed at 180 degrees from each other in an otherwise uniformly illuminated arena (*30*). The two stripes are unreachable by the fly because the walking arena is separated from the stripes by a trench of water, inducing the fly to walk back and forth between the two stripes for the duration of the assay. A major advantage of open world assays like Buridan’s paradigm is the ease of repeated measurement of a large number of behavioral parameters on freely walking individuals. This assay is named after a medieval philosophical paradox meant to highlight the importance of intrinsic bias when external conditions are completely symmetrical. Interestingly, the problem of how to resolve identical options has been proposed as one of the advantages of a non-deterministic and noisy brain (*31*).

Using this behavioral paradigm, we find that flies show highly idiosyncratic responses that are very stable over a long period of time. In particular, the width of the path that a fly walks between the two stripes, a parameter we call “absolute stripe deviation”, is a unique and stable feature of a given fly that shows a normal distribution of variability within the population. We show that behavioral individuality of stripe deviation is non-heritable and is not reduced through inbreeding. Using unbiased anatomy-to-behavior correlation mapping, we find that the degree in left/right DCN wiring asymmetry is a robust predictor of behavioral performance of an individual fly and its variance across the population. The more asymmetric the DCN wiring pattern, the narrower the path a fly walks between the two stripes. DCN activity is necessary for this correlation, while artificially inducing asymmetry in the visual system is sufficient to change the response of an individual. This establishes a causal link between variability in the development of the brain and the emergence of individuality of animal behavior.

## Results

While analyzing object orientation responses in wildtype *Canton S* (*CS*) flies using Buridan’s paradigm (Fig. 1 A and Movies S1-3) we noted significant degree of inter-individual variability in their trajectories. Given that males and females display dimorphic behavioral traits (*32*) and are genetically different from each other, we first tested whether this variation was principally due to sex differences. However, at the population level males and females displayed similar responses towards the high contrast stripes, as shown by the occupancy heatmaps and individuals with equally variable responses were found in both sexes (Fig. 1 B-C). Next, we used a simple and robust parameter called “absolute stripe deviation” which shows how much a fly deviates from an idealized narrowest possible path between the stripes. Thus, a low stripe deviation score indicates a fly that walks a narrow and straight path between the two stripes, while a high stripe deviation score indicates a fly that walks a broad and meandering path between the two stripes. We found that males and females display a similar degree of inter-individual variation in stripe fixation (Fig. 1 D). Therefore, we combined the responses of the *CS* population in one histogram displaying the variability for absolute stripe deviation (Fig. 1 E) and continued our studies with combined male and female populations.

**Fig.1.**
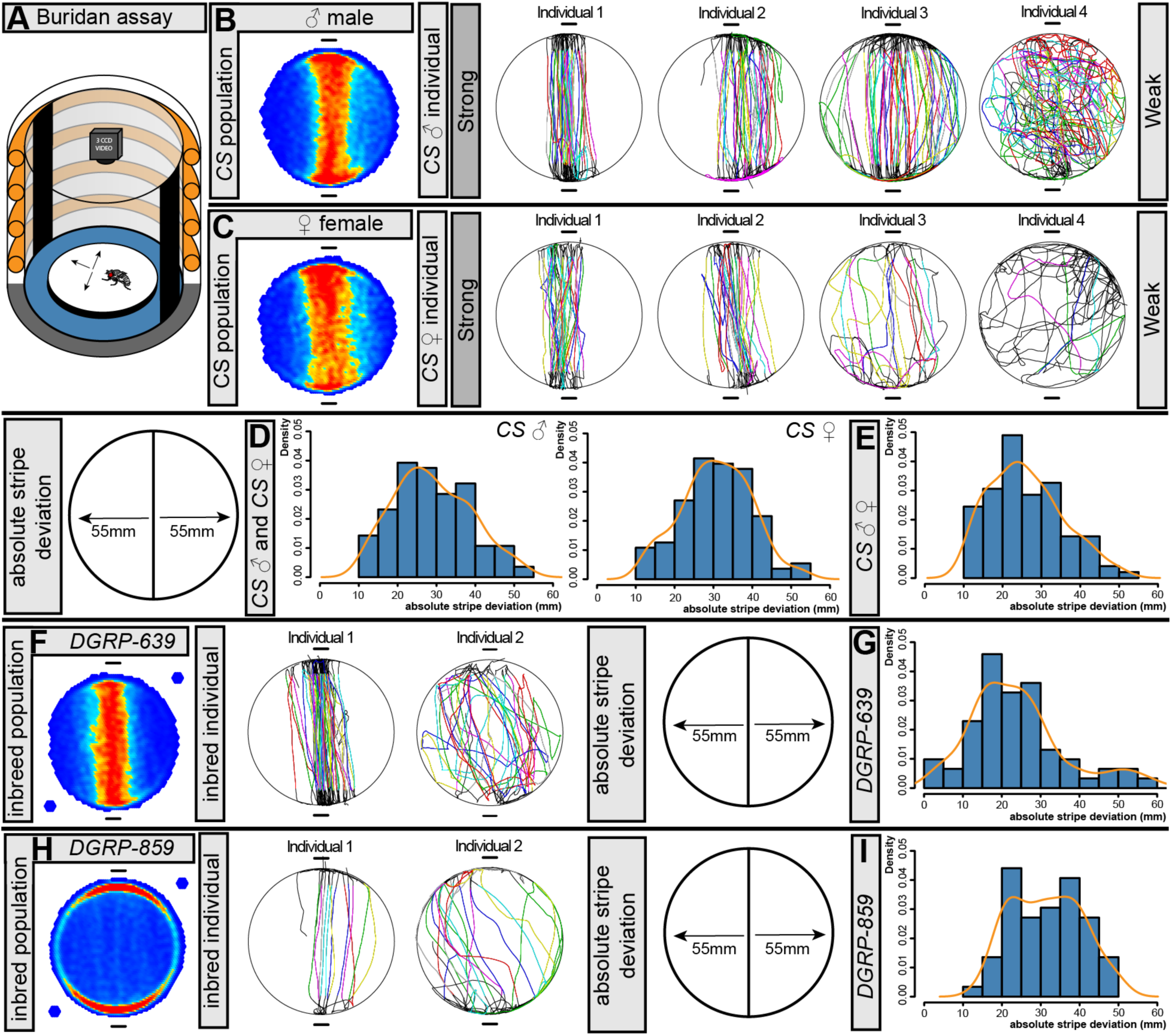
Individual variation of Drosophila stripe responses is independent of gender or genetic diversity. A) Drosophila object orientation responses are measured in an evenly illuminated Buridan’s paradigm arena. The fly is located on a round platform. The two vertical black stripes are the only visual cues. All fly tracks are recorded with a high definition digital camera. B) Male CS wildtype flies (N = 50) showed on the population level (shown in a heatmap) a clear object orientation response towards the stripes that are located at the top and bottom. The categorization of individual responses into strong and weak object orientation responses shows the entire repertoire of responses. C) Female CS wildtype flies (N=48) showed on the population level (heatmap) the same object orientation response as their male counterparts. The same was also true for the individual variation that we observed also for female flies. D) The histograms for absolute stripe deviation shows that CS male (N = 50) and female (N = 48) flies show the same range of individual responses. The distributions are statistically identical (Tukey Test, p-value = 0.1). E) The histogram shows the cumulative stripe deviation for CS male and female flies (N = 98). F) DGRP-639 flies (N = 61) showed on the population level (shown in a heatmap) an object orientation response towards the stripes that exceeds even the response of CS (Tukey Test, p-value = 0.01). Two examples of individual responses show the individual differences. G) The histogram for absolute stripe deviation shows that DGRP-639 flies (N = 61) exceed the variability of CS flies (F-Test, p-value<0.001). The distribution is shifted towards lower stripe deviations. H) DGRP-859 flies (N = 59) showed on the population level (heatmap) a weak object orientation response towards the stripes. The main population response is edge behavior. Two examples of individual responses show prevalent individual differences that include also individuals with strong object orientation responses. I) The histograms for absolute stripe deviation show that DGRP-859 flies (N = 59) show comparable variability as CS flies. The distribution is shifted towards higher stripe deviations.

### Individual variability of object orientation responses is independent of genetic diversity

Next, we asked if the behavioral variability correlates with genetic diversity. If genetic diversity results in behavioral variability, then strains with low genetic diversity should show reduced object orientation variability. To test this idea, we compared strains with high genetic diversity to strains with low diversity. First, we screened a subset (N=10) of the *Drosophila* genomic reference panel (DGRP) for strains that showed either very strong or very weak object orientation responses at the population level. DGRP lines are fully sequenced and they have been inbred for 20 generations, which should make all individuals of one strain as genetically homogenous as it is compatible with viability (*33*). We identified two strains with opposing behavioral phenotypes at the population level. *DGRP-639* showed near wildtype absolute stripe deviation (Fig. 1 F-G), while *DGRP-859* showed strongly increased stripe deviation (Fig. 1 G-I). Importantly, these population level differences were not confined to the stripe deviation index as statistical analysis of eight representative behavioral parameters showed that the outbred *CS* control strain differed significantly in the mean from the inbred lines (Fig. S1 A), showing that these were truly behaviorally different populations. Next, we asked if the inbred DGRP lines showed a reduction in behavioral variability along with their reduced genetic diversity. We find that the degree of individual variation of object orientation responses, as indicated by the distribution of the stripe deviation index, was not reduced (Fig. 1 G, I). In fact, if anything, the *DGRP-639,* which as a population has reduced stripe deviation, showed increased behavioral variability (Fig. 1 G, Fig. S1 A). Therefore, reduction of genetic diversity did not reduce phenotypic behavioral variation.

### Individual object orientation responses are non-heritable

We next wanted to know whether a specific genotype in an individual is associated with reproducible behavior of that individual. If that would be the case, we should be able to breed specific alleles by choosing parental animals with certain behavioral traits. From a *CS* population of 47 male and 37 virgin female flies, we selected and mated the three pairs with the lowest and highest stripe deviation scores, respectively (Fig. 2 A). Object orientation responses in the offspring of these pairs were measured in the Buridan arena (Fig. 2 B, C). We find no differences between the two sets of offspring in stripe deviation scores as well as six other parameters tested (Fig. 2S A). The same was true for the offspring of a single pair with low and high stripe deviation (Fig. 2S B-C). More importantly, the individual variation of stripe deviation scores of the two sets of offspring matched each other, as shown in the histograms, and matched the original variability of the parental *CS* population. Therefore, an individual’s behavioral profile is non-heritable and offspring of similar parents recreate the individual variability profile of the starting population.

**Fig.2.**
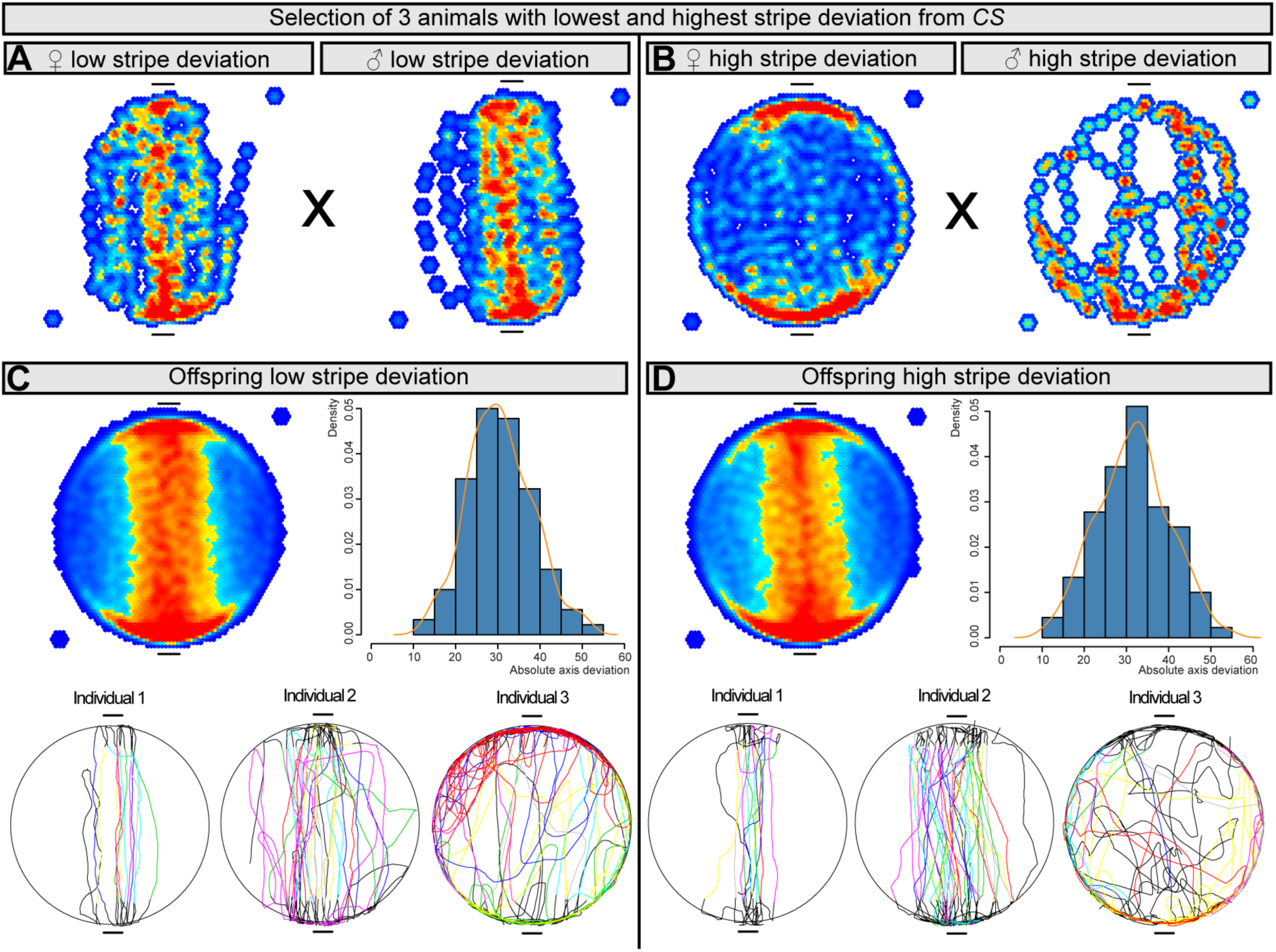
Individual variation is independent of genetic selection. A-B) In a selection experiment the three best (A) and worst couples (B) for absolute stripe deviation were chosen from a population of 47 CS males and 37 virgin CS females. The heatmaps in the top row show from left to right: 1. the three virgin females and 2. males with the lowest stripe deviation, 3. the three virgin females and 4. males with the highest stripe deviation. C-D) The offspring of these two populations is shown in the middle row separated by a black stripe. Neither the heatmaps or the histograms make any of two populations (N = 180, 180) different from each other. The statistical analysis (two-way ANOVA and Tukey HSD as post hoc test) for stripe deviation shows with a p-value of 0.22 no statistical difference between the two populations. The same is true for most of the other path parameters we looked at. In the bottom row we show examples of individuals representing the range of variability in both populations. No difference was found for variability between the two populations.

### Individual object orientation responses are stable over time

A specific behavioral profile may not be heritable for two reasons. First, because it is driven by current state modulations and is thus not a stable trait of an individual. Second, because it is a stable trait driven by intrinsic but non-heritable mechanisms. To distinguish these possibilities, we asked if the object orientation responses of an individual are stable over extended periods of time. We tested the same individual *CS* flies once every other day for three days, in the Buridan arena. We find that an individual’s behavior is virtually identical over the three trials for animals with low (Fig. 3 A) as well as high (Fig. 3 B) stripe deviation scores. Statistical analysis of absolute stripe deviation showed that the individual responses of *CS* flies on different days were strongly and highly significantly correlated (Pearson correlation coefficients ranging from 0.74-0.77, Fig. 3 C). Furthermore, the same was true for path details like left or right shifted angles (Fig. 3 A and Fig. S3 A for statistical analysis, including angle parameters). Similar results were obtained for many other behavioral parameters including: distance, full walks, meander, absolute horizon deviation, absolute angle deviation, angle deviation and center deviation (Fig. S3 C). The stability of behavioral responses over several days argued strongly against current state modulations and in favor of individual properties. Next, we asked how temporally stable object orientation responses are by testing four-week-old flies repeatedly in our arena. Aged animals, exactly like their young counterparts showed highly stable individual responses (Fig. S3 D). Finally, we asked whether reduced genetic diversity might impact behavioral stability. We performed repeated testing of *DGRP-639* and *DGRP-859* individual flies and found that both inbred strains showed temporally stable individual stripe deviation responses (Fig. 3 D-E; Fig. S3 A-B), as well as many of the other behavioural parameters for both DGRP lines (Fig. S3 C).

**Fig.3.**
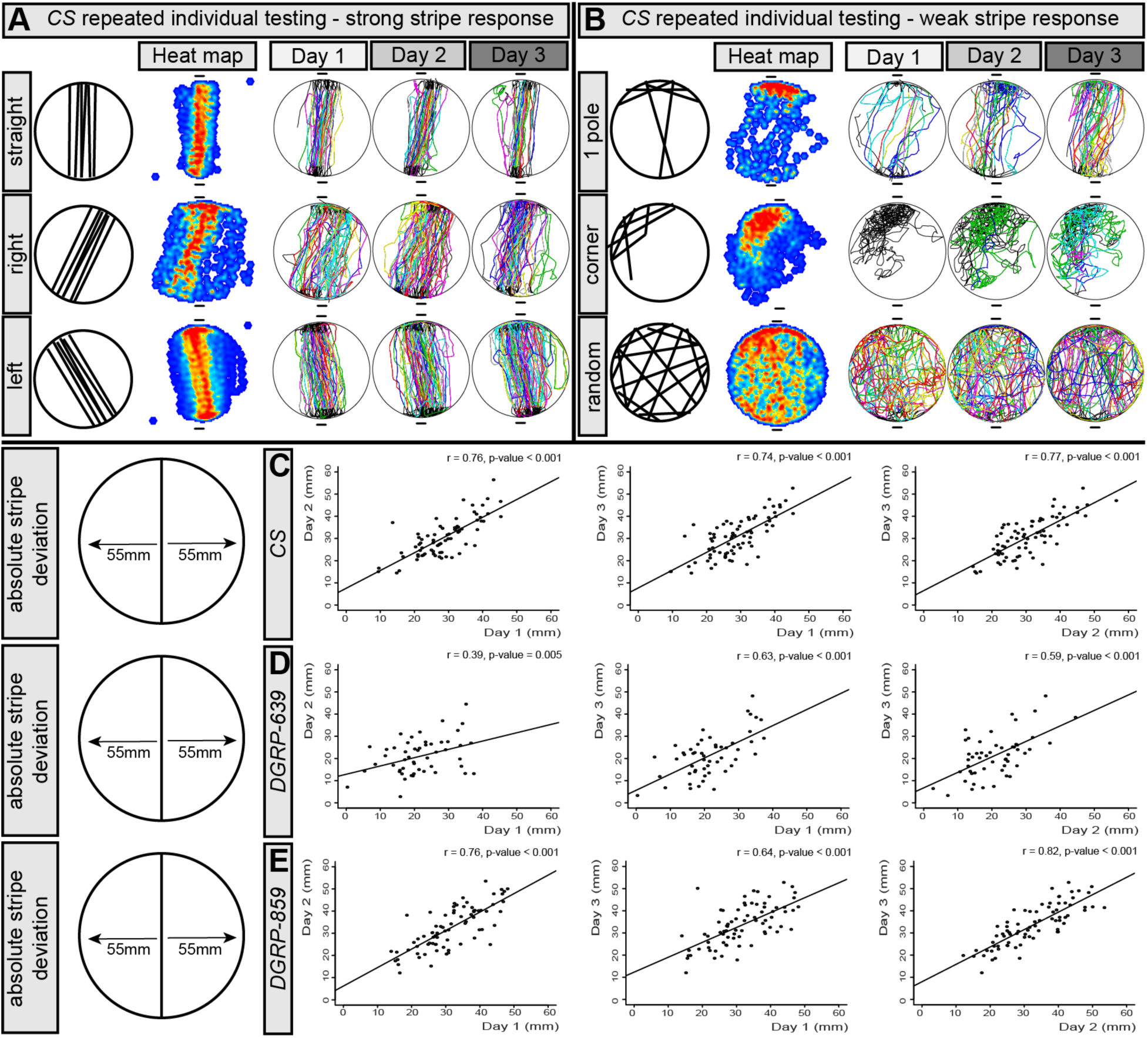
Individual variation of Drosophila object orientation responses is stable over time. Adult CS flies (N = 74) were repeatedly tested over the duration of three days. The flies showed remarkably stability in their responses irrespective of whether they responded strongly or weakly to the visual cue, or even showed no response towards the visual cue at all. A) Three different examples of CS flies with strong stripe fixation. The heatmap and the individual tracks for the three days show highly persistent behavior. Even the angle towards the stripes is highly conserved throughout the three days. The upper row animal shows no apparent angle, the middle row is right shifted, and the bottom row is left shifted. B) Three different examples of CS flies with weak stripe fixation. The positional preferences are highly conserved between the different days. The upper two rows show animals with specific local preferences and the lower one shows a random walk. C) Statistical analysis for absolute stripe deviation shows that the CS responses of the different days are strongly correlated. The Pearson correlation coefficient for Day 1 vs. Day 2 is 0.76 with a p-value < 0.001. For Day 1 vs. Day 3 the correlation coefficient is 0.76 with a p-value < 0.001. For Day 2 vs. Day 3 the correlation coefficient is 0.76 with a p-value < 0.001. D) Similar to the CS data, the responses for DGRP-639 were moderately to strongly correlated (N = 52, animals with extremely low path length were removed, see Fig. S3 A for examples). The Pearson correlation coefficient for Day 1 vs. Day 2 is 0.39 with a p-value=0.0047. For Day 1 vs. Day 3 the correlation coefficient is 0.63 with a p-value<0.001. For Day 2 vs. Day 3 the correlation coefficient is 0.59 with a p-value < 0.001. E) The correlation between days for DGRP-859 (N = 76) even exceeds the data for CS. The Pearson correlation coefficient for Day 1 vs. Day 2 is 0.76 with a p-value<0.001. For Day 1 vs. Day 3 the correlation coefficient is 0.64 with a p-value<0.001. For Day 2 vs. Day 3 the correlation coefficient is 0.82 with a p-value < 0.001. Table 1. Start this caption with a short description of your table. Format tables using the Word Table commands and structures. Do not create tables using spaces or tab characters.

Altogether, our behavioral analyses suggest that individual variability in object orientation responses is an intrinsic, non-heritable, temporally stable trait that is independent of sex and genetic background and that is not eliminated by reduced genetic diversity. What in the brain might underlie such individuality in visual behavior?

### The DCNs are a highly variable set of commissural visual interneurons

In a classic object orientation paper Bülthoff (*34*) suggested, based on earlier work by Zimmermann and Götz (*35, 36*), that object position processing in *Drosophila* and other insects (*37*) might in part rely on what was referred to as qualitative asymmetry between front-to-back and back-to-front motion. This simply refers to the fact that as a fly turns towards an object, this object moves from front-to-back on one retina, and from back to front on the other retina. Asymmetry of this percept would allow the fly to better center the object in the frontal part of its visual field. However, direct evidence for this notion remains lacking, and it is unclear where the source of such visual system asymmetry would be, especially that the sizes of the left and right eyes of the same fly, though not identical, are nonetheless very highly correlated (*38*). In 1986, Heisenberg and colleagues suggested that binocular interactions, perhaps through commissural interneurons further upstream in the visual system may be required for object orientation in the frontal visual field (*39*). Interestingly, the DCNs described above precisely match this predicted circuit. DCNs, whose function has thus far remained unknown show a highly variable wiring diagram between individuals and between the left and right sides of the same individual, due to intrinsically stochastic developmental wiring mechanisms (*21, 22*). DCN soma are located on the dorsal side of the fly visual center called the optic lobe. Each DCN projects a single ventral neurite that splits into an ipsilateral dendrite and a contralateral axon. On the contralateral side the axon terminates in either the lobula, thus defining the neuron as a L-DCN, or the medulla for the M-DCNs (Fig. 4 A). The DCNs show variability on all three levels: the number of cell bodies, the number of axons innervating the lobula and the number of axons innervating the medulla (*21*). All three levels of variability are observed between individuals as well as between the left and right hemispheres of the same individual.

**Fig.4.**
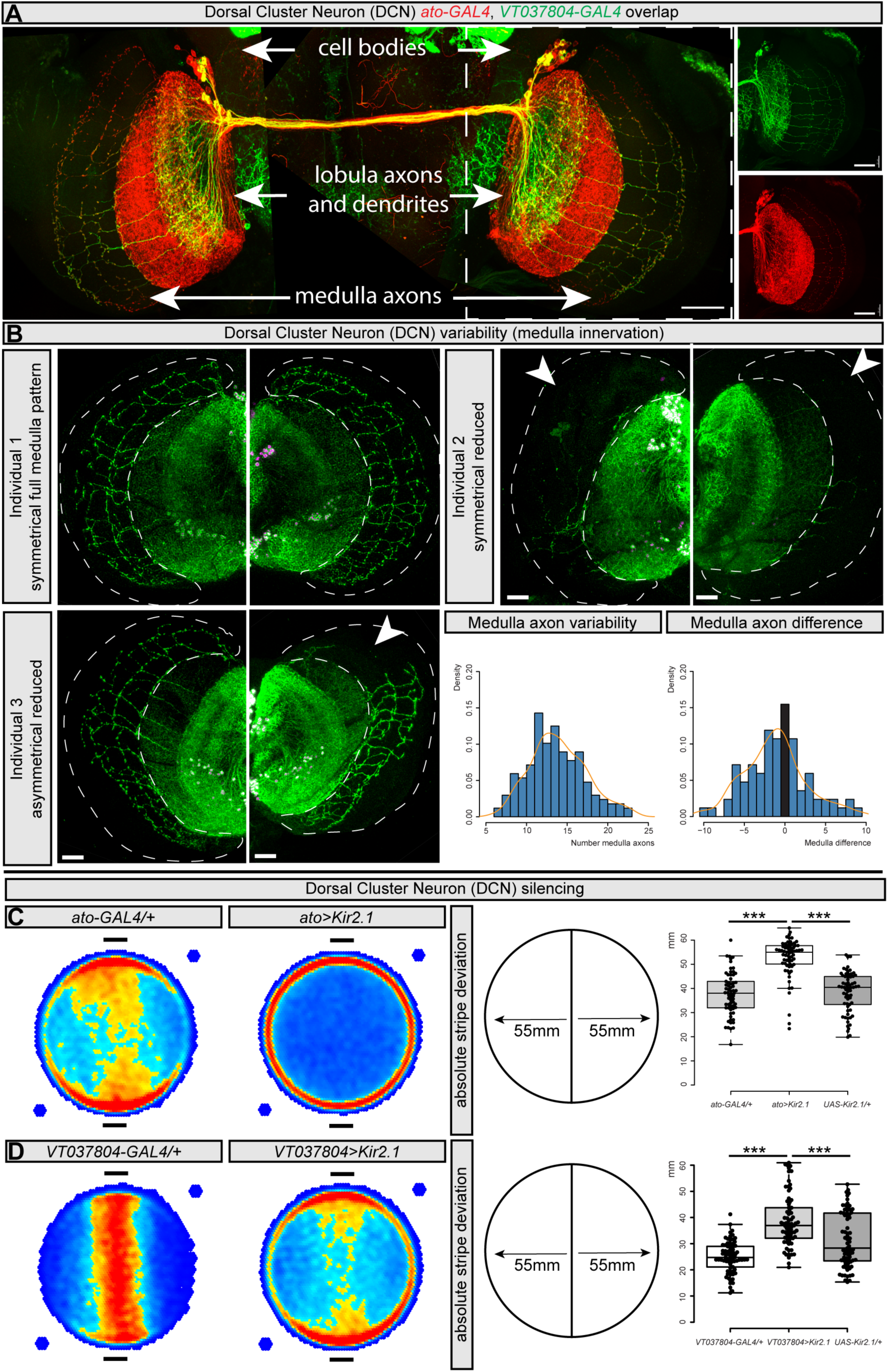
Normal stripe responses require Dorsal Cluster Neuron function. A) The Dorsal Cluster Neurons (DCN) are commissural neurons in the visual system of the fly. The DCNs have dorsally located cell bodies that sends out an ipsilateral dendrite and a contralateral axon. This axon either innervates the visual neuropil lobula or medulla. Two independent driver lines are shown for the DCN neurons: ato-lexA (red, lexAOP-myr-tdTomato) marks all DCNs while VT037804-GAL4 (green, UAS-myr-GFP) marks only the M-DCN neurons that innervate the medulla. B) The DCNs show high variability in their axonal branching pattern as shown for three individual brains. Green shows ato>mCD8-GFP and magenta ato>red-Stinger. The medulla is outlined by a dashed line and missing axonal innervation is marked by an arrowhead. The first individual shows a full DCN medulla innervation pattern in both brain hemispheres, the second individual has in both brain hemispheres medulla innervation missing, the third individual has one complete hemisphere and one hemisphere with missing medulla innervation. Statistical analysis shows that the number of medulla axon branches ranges from 6 to 23 axons with a mean of 13.99. The medulla asymmetry ranges from 0 to 10 axons with a mean of 2.98. C) DCN neuron silencing leads to an almost complete loss of the stripe fixation response. The heatmap of the control population of ato-GAL4/+ flies show a normal response in the two-stripe arena. This is entirely lost upon silencing of DCN neurons in ato>Kir2.1 animals. Statistical analysis (N=57-63 Set) of the absolute stripe deviation shows that the ato>Kir2.1 animals show significant higher stripe deviation than the controls (two-way ANOVA and Tukey HSD as post hoc test p-value<0.001). Higher stripe deviation means that the animals fixate the stripes less. D) Similar results are obtained by M-DCN neuron silencing with VT037804-GAL4. The heatmap of the control population of VT037804-GAL4/+ flies shows normal fixation in the two-stripe arena. This is entirely lost upon silencing of DCN neurons in VT037804>Kir2.1 animals. Statistical analysis (N=72 Set) of the absolute stripe deviation shows that the VT037804>Kir2.1 animals show significant higher stripe deviation than the controls (p-value<0.001). Scale bars correspond to 20 μm.

We first extended the quantitative analysis of DCN wiring variability using an automatic neuronal reconstruction method on a much larger number of individuals than previously analyzed (N = 103). Our data revealed that we had in fact previously underestimated DCN variability (*21*). Specifically, we found that the number of DCNs varies from 22-68 cells, with a range of 11-55 L-DCNs and 6-23 M-DCNs (Fig. 4 B; Fig. 4S A). In addition to the obvious asymmetries in the number of neurons the majority of which are L-DCNs, we observe a distribution of variation in medulla targeting asymmetry by M-DCNs (Fig. 4 B, histogram distributions; Fig. 4S A). The distribution of all DCN asymmetries showed a peak of low asymmetries, while extreme asymmetries were present but rare. Importantly, this extended analysis confirmed the previous observation that the number of axons in the medulla is not correlated to the total number of DCNs that give rise to them (Fig 4S A). In other words, the number of DCNs in the right hemisphere does not predict the number of DCN axons in the left medulla and *vice versa*. Finally, 3D reconstruction shows that M-DCN axons terminate specifically in the posterior medulla (Movies S4-6), where visual columns form the anterior (i.e. frontal) visual field are located, and the DCN wiring pattern in the medulla does not change in the adult (Movie S7). In summary, the DCNs innervate the frontal visual columns in the medulla in a fashion that shows a variable degree of asymmetry between individuals which is stable over time in any given individual.

### Individual variability in DCN wiring asymmetry drives individual variability in object orientation behavior

The data above suggest that DCNs represent an ideal candidate for an intrinsically asymmetric population of contralateral higher order interneurons that may mediate binocular interactions relevant to the anterior visual field (*39*). We therefore hypothesized that the DCNs represent a circuit that explains how binocular asymmetries in the visual system regulate object orientation responses. To test this hypothesis, we first asked whether the DCNs were required for object orientation in the Buridan assay (*29*). Silencing either all DCNs, or only M-DCNs, by expressing either the potassium ion channel Kir2.1 (*40*) or the tetanus toxin light chain (TNT) (*41*) resulted in a strong disruption of object orientation behavior. Specifically, DCN silenced flies showed a strong tendency to approach the two stripes from the extreme edges of the arena, as opposed to through the middle, as revealed by the change in the occupancy heat maps and a significant increase in the stripe deviation index (Fig. 4 C, D and Fig. 4S B, C).

Next, we queried the relationship between individual variability in object orientation behavior and individual variability in DCN cell numbers and wiring diagram. To this end, we expressed nuclear and membrane markers in DCNs and quantitatively measured individual object orientation behavior of these flies (N = 103), followed by high-resolution confocal imaging and semi-automatic neuronal reconstructions of each individual, and finally, an unbiased correlation analysis between 36 key behavioral parameters and 37 prominent anatomical features of DCNs in each individual (Fig. 5 A, Fig. 5S A-C). We found that left/right asymmetry in medulla innervation by DCNs strongly and specifically correlated with an individual’s stripe deviation index (Fig. 5 B, r = −0.67) and other inter-dependent parameters, such as absolute angle deviation and center deviation, but not unrelated parameters such as total distance or the number of full walks between stripes (Fig. S5 A). Individuals with high M-DCN asymmetry tend to have a low stripe deviation index (i.e. walk a narrow path between stripes), while individuals with symmetric M-DCN have a high stripe deviation index (Fig. 5 C).

**Fig.5.**
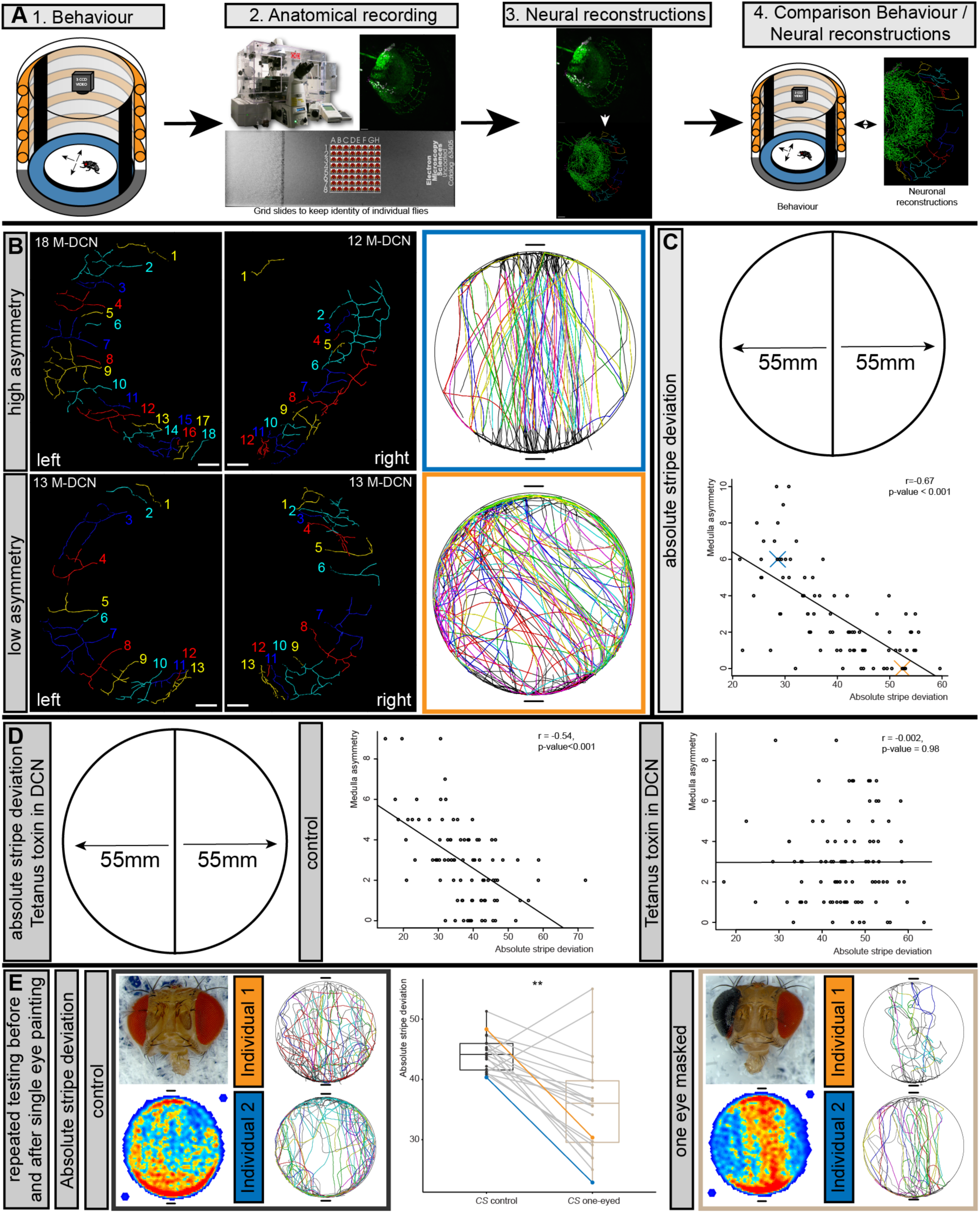
Individual variation correlates with anatomical brain asymmetry. A) To correlate behavioral variation with brain anatomical variation we first tested flies behaviorally in the Buridan arena. Afterwards we recorded the brain anatomy keeping individual information intact. After computational analysis of fly tracks and automated neuronal reconstruction of the brain anatomy, we performed a cross-correlational analysis of both data sets. B) The most striking correlation between behavior and anatomy was that animals with high asymmetry performed stronger than animals with lower asymmetry of DCN axonal projections. The colored numbers show the reconstructed medulla axons in each brain hemisphere. C) Statistical analysis (N = 103) shows that the medulla asymmetry correlates strongly (r = 0.67, p-value < 0.001) with the absolute stripe deviation. The blue and orange crosses mark the position of the blue and orange individuals shown in B). D) Dorsal cluster neuron silencing with Tetanus Toxin (TNT, N = 92) results in the loss of this correlation (r = 0.002, p-value = 0.98; control: N = 89, r = 0.54, p-value < 0.001). E) Wildtype CS flies were repeatedly tested for before and after blinding a single eye, resulting in an asymmetric visual input. The data shown represents only animals with an absolute stripe deviation above 40 that should have a symmetrical DCN pattern (N = 20). Upon blinding a single eye, the path as measured by absolute stripe deviation significantly improves (paired Wilcoxon Test, p-value = 0.002).

If DCN wiring asymmetry is a functional driver of individual object orientation behavior, then silencing DCN activity should abolish the correlation between anatomy and behavior in individuals. To test this, we expressed TNT in DCNs labeled with GFP and repeated the individual behavioral-anatomy correlation analyses as described above. We observed that in animals that express TNT in DCN neurons, no significant correlation exists between M-DCN asymmetry and absolute Stripe deviation (Fig. 5 D, r = −0.002), demonstrating a requirement for DCN activity for a link between wiring asymmetry and behavior in individuals.

### Visual asymmetry determines object orientation in individuals

Our data show that under normal conditions intrinsic, non-heritable, developmental variation in DCN wiring asymmetry is necessary for creating variability in object orientation behavior across individuals. This supports Götz’s original hypothesis that object orientation depends on asymmetry in the processing of visual input. To directly test if generating asymmetry anywhere in visual information is sufficient to change an individual’s behavior, we tested 79 wildtype *CS* flies in the Buridan arena. Next, we selected the 20 flies with stripe fixation indices above 40 – thus those that tend to have lower asymmetry in DCN wiring – and then performed monocular deprivation and tested them again. We find that monocular deprivation resulted in a significant reduction of the stripe deviation index in these flies (Fig. 5 E). Remarkably, this was also true for the entire population (Fig. 5S D). Therefore, variability in the asymmetry of visual processing causally contributes to the behavioral variability in object orientation behavior.

## Discussion

The question for the origins of individual behavioral variation a central open question in the neurosciences and psychology. Obviously both heritable and environmental factors shape any given individual, yet mechanisms that explain behavioral individual variation in most cases remain elusive. The discovery of stable individual traits in non-human vertebrates (*20*) and invertebrate species facilitated research in numerous species on behavioral variation (*14, 17*). In recent years, due to experimental advantages, invertebrate model systems like the fruit fly *Drosophila melanogaster* and the nematode *Caenorhabditis elegans* have become increasingly popular to answer important questions about the origins of individual behavioral variability (*15, 16, 19, 42*). These seminal papers clearly demonstrate that individuality can be found in genetic invertebrate model organisms and they offer both genetic (*15, 16, 42*) and neuromodulatory (*15, 19, 42*) explanations for the observed idiosyncrasies.

The work presented here demonstrates an entirely new mechanism to explain aspects of individual behavioral variation through stochastic variability in nervous system development. We had previously reported that significant wiring variability in a population of higher order visual system neurons called the DCNs arises through fundamentally stochastic cell-cell interaction mechanisms such as lateral inhibition and cell surface receptor recycling during filopodial growth and retraction (*21, 22*). This study further shows that these mechanisms also give rise to significant intra-individual variation in the shape of a broad distribution of left/right asymmetry in the innervation of the visual areas by contralateral DCN axons. Such extensive individual variability in neural circuit development raises the question of whether individual variation in circuit morphology has implications for individual variation in visually guided innate behavior. We addressed this question using the Buridan paradigm (*29*), a robust object orientation assay that allows repeated testing of the same individual. Our data show that flies display temporally stable, non-heritable, individual behavioral differences in object responses. When considered in the context of previous reports showing individuality in similar to locomotor handedness (*16, 42*) and phototactic (*15*) behavior in flies, our work shows that flies have innate individuality traits across a complex range of visually guided behaviors. The amenability of the relatively complex *Drosophila* brain to multiscale analysis, from the molecular to the behavioral, at single animal resolution makes it an ideal model for understanding the emergence of individuality at each of these scales.

Our work demonstrates that variability in neuronal wiring arising during development strongly correlates with behavioral individuality. Specifically, using an unbiased approach in which object orientation behavioral parameters were correlated with DCN wiring morphology, we show that the variation in left/right asymmetry across individuals in a population explains the variation in object orientation responses. In this regard, two observations merit further comment. First, unbiased analysis revealed a highly specific correlation between the shape of the path a walking fly takes towards an object and the asymmetry of DCN innervation of a particular visual area called the posterior medulla, through which information form the frontal visual field flows. Individuals with high asymmetry show much stronger object orientation responses than animals with high symmetry. Second, silencing DCN activity did not abolish object responses per se, but rather completely abolished the correlation between wiring variation and behavioral variation, meaning that the anatomical asymmetry is functionally relevant. Given the very little asymmetry observed in eye size in *Drosophila* lab strains, our work suggests that in the fly visual system asymmetries relevant to object orientation arise in the brain and not in the periphery. To query whether peripheral asymmetry has the same effect, we blocked a single eye of animals with weak orientation responses and found that this strongly enhanced their object orientation, meaning that asymmetry in visual information is in and of itself sufficient for improving object orientation. This is important because classic work in *Drosophila* visual behavioral neuroscience lead to the proposal that asymmetry in visual information processing influences object responses. Where such functional asymmetry lay and how it might arise has, until now, remained unclear. Independently, the study of object responses in motion blind mutants led Heisenberg and colleagues to propose a hypothetical contralateral circuit dedicated to object responses in the frontal visual field (*30, 34-36, 39*). We therefore propose that the DCNs are the neurons that explain both of these observations: a contralateral asymmetric visual circuit that regulates object orientation in the frontal visual field. Future work will reveal the exact physiological consequences of morphological asymmetry, such as whether wring asymmetry induces timing differences as in auditory navigation (*43*) or whether the absolute differences are simply summed up.

Here we show that inherently stochastic cellular mechanisms lead to probabilistic wiring diagrams across a population which in turn underlies significant individual variability in behavior in an invertebrate model. This is consistent with the fact that variations in fluctuating morphological asymmetry arise during development even in genetically identical individuals of parthenogenic species raised in the same microenvironment (*44*). Studies in humans clearly show similar correlations between variations in brain morphology and individual variations in behavior and personality (*27, 45*). More recently, some studies have begun to focus on those anatomical features that can be definitively traced back to developmental events. Work focused on differences in reading capabilities shows that continuous versus interrupted morphology of the human sulcus in the visual word form area, which arises during fetal development, predicts reading skills in adults (*46*). This, combined with differences in brain anatomy between identical human twins, strongly suggests that stochastic developmental variation in neural network formation is, in addition to genetic differences and environmental factors, a determining factor of individual behavioral variation and personality. The amenability of invertebrate models to highly detailed multiscale analysis, from the molecular to the behavioral, at single animal resolution of the causal links between genes, development and the environment in generating personality traits holds great promise for further breakthrough discoveries in the field, that the history of science shows will likely translate to human biology.

## Acknowledgments

We thank the Bloomington stock center (NIH P40OD018537) and Gilad Barnea for providing flies used in this study. We thank Bruno van Swinderen and all members of the Hassan, Hiesinger and Wernet lab for support and providing valuable comments in the course of this project. We further would like to thank the VIB Bio Imaging Core for support

## Author contributions

G.A.L. and B.A.H conceived the study, designed the experiments and wrote the manuscript. G.A.L. conducted all behavioral experiments, immunohistochemistry and all data analysis. M.A., S.D. and L.H. helped with the neuronal reconstructions. G.L. and R.K.E provided technical expertise and R.K.E wrote the python analysis software. L.M.F and A.D.S shared data before publication and provided technical expertise. M.W. and P.R.H. provided expertise and equipment and helped writing the manuscript

## Supplementary Materials for A neurodevelopmental origin of behavioral individuality

### Materials and Methods

#### Fly stocks and rearing

Fly rearing and immunohistochemistry was performed according to standard procedures (*1*) (see extended materials and methods). All experimental animals were kept under non-crowded conditions.

The following fly stocks were used: *Canton S* (*CS*) as outbred wildtype stock *DGRP-639* (Bloomington stock number 25199) and *DGRP-859* (Bloomington stock number 25210) (*2*) as inbred wild-type stocks. GAL4s: *ato(14A)-GAL4* (*3*), *VT037804-GAL4*, UAS: *UAS-mCD8-GFP*(*4*), *UAS-redStinger* (*5*), *UAS-Kir2.1* (*6*) *UAS-TNT* (*7*), *UAS-trans-Tango* (*8*).

Stocks and experiments were reared on a standard cornmeal/agar diet (7.5g Agar, 64g cornmeal, 160g yeast, 85.5ml sugar cane syrup, 8.5ml ethanol, 0.51g Nipagin and 2.5ml propionic acid in 1L of water). Experimental animals were reared in low densities (two females per vial) with frequent food changes to prevent crowding effects at 25°C in a 12/12-hour light/dark regime at 60% relative humidity. Only the first two days of offspring per vial were used for any experiment.

#### Immunohistochemistry

*Drosophila* brains were dissected in PBS and fixed in 3.7% formaldehyde for 20 min. Subsequent washes and incubations were done in PBS with 0.2% Triton. Tissues were incubated overnight with primary antibody at 4°C, followed by a 2 hour incubation with secondary antibodies at room temperature the next day. The following antibodies were used: goat α-green fluorescent protein (GFP) (Abcam, 1:2000) and rat α-N-Cadherin (DSHB,1:50)(*9*), FITC-, Cy3-, and Cy5-conjugated secondary antibodies were obtained from Jackson Immunolabs and used at 1:200 (1:100 for the Cy5-conjugated antibody). Fluorescent preparations were mounted in Vectashield (Vector Labs). Images were acquired using Leica SP8, SPE and a Nikon TI2000 AIR confocal microscopes.

#### Buridan’s paradigm

Buridan experiments were performed according to published protocols(*10*), except that the temperature on the platform was 25°C and experiments were started 4-7 days after eclosion and at least 2 full days (48 hours) were waited after CO_2_ anesthesia.

The Buridan arena consisted of a round platform of 117 mm in diameter, surrounded by a water-filled moat. The arena was placed into a uniformly illuminated white cylinder (Fig. 1 A). The setup was illuminated with four circular fluorescent tubes (Osram, L 40w, 640C circular cool white) powered by an Osram Quicktronic QT-M 1×26–42. The four fluorescent tubes were located outside of a cylindrical diffuser (DeBanier, Belgium, 2090051, Kalk transparent, 180g, white) positioned 147.5 mm from the arena center. The temperature on the platform during the experiment was 25°C. Except in experiments where flies were walking without visual stimuli, 30 mm wide stripes of black cardboard were placed on the inside of the diffuser. The retinal size of the stripes depended on the position of the fly on the platform and ranged from 8.4° to 19.6° in width (11.7° in the center of the platform).

Fly tracks were analyzed using CeTrAn(*10*) and custom written python code (available upon request).

The following parameters were analyzed:

Speed: Median speed was calculated by dividing the distance travelled by time. Speeds exceeding 50 mm/s were considered as a fly jumping and were excluded in the speed calculation.

Distance: All movements were added up over the whole experiment to result in the total distance traveled by each fly (in mm).

Turning angle: The median turning angle was calculated for each fly (in degrees).

Meander: a measure of the tortuosity of the trajectories. It was calculated by dividing the turning angle by the instantaneous speed. The median was calculated for each fly (in degrees × s/mm).

Centrophobism (moving and stationary): The circular arena was divided into a smaller disk and an outer ring of equal surface (taking a disk of a radius √2 times smaller than the platform radius). The software determined the time spent in each subdivision, treating data points while the animal is in motion independently from data points where the animal was stationary. The centrophobism indices for moving and for stationary (respectively) were then calculated as the difference between the number of data points outside and inside of the center area, divided by the sum of the two numbers. Therefore, an index of 1 would mean that the fly spent the entire experiment in the outer area, while a −1 would mean that the fly spent the entire experiment in the center and 0 denotes an equal distribution between outside and inside areas of the platform.

Absolute angle deviation: This metric corresponds to the angle between the velocity vector and a vector pointing from the fly position toward the center of the front stripe. For each displacement, the vectors going from the fly position toward both stripes (situated at p (0.0 +/-146.5 mm) in the new coordinates centered in the platform center) are calculated and the respective angles between the velocity vector and each of those vectors are measured. Finally, the smaller of the two angles is chosen as output. The median of all deviation angles is reported for each fly (in degrees). Smaller values then correspond to a path directed toward the stripes.

Full walks: This metric corresponds to the number of times the fly walked from one stripe to the other (closer than 80% of the platform radius toward the stripe, Fig. 3). The software detects when the fly enters one of the two areas and increments the count by one when it enters the opposite area. This process is reiterated until the trajectory ends.

Activity metrics: From the speed profile of the trajectory (instantaneous speed over experimental time), there are different ways to determine an activity pattern. Our first computation (time-threshold: indices labeled with (TT)) considers every movement as activity and every absence of movement lasting longer than 1 s as a pause (shorter periods of rest are considered as active periods). Changing this threshold from 1 to 0.5 or 1.5 s had little effect on the results (data not shown), such that we arbitrarily chose 1 s as standard.

For both activity computations, we calculated for each fly the total activity time (in seconds), the number and median duration of the pauses and the median duration of bouts of activity (in seconds). For the median duration of activity (TT), we made a second calculation considering only activity bouts leading to larger displacement (>1 cm).

Custom written positional metrics (Python, Python Software Foundation):

Angle deviation: This metric was calculated similar to absolute angle deviation with the addition that the directionality of the angles mattered and a deviation from the right part of the arena (in accordance to the directionality of the vector) gave a positive angle, while on the left a negative angle was recorded.

Stripe deviation: This metric was calculated as the deviation of the position away from the imaginary line through the two stripes. Both the directional stripe deviation and the absolute stripe deviation were calculated. In the directional stripe deviation, the right part gave positive and the left gave negative values. For absolute stripe deviation, it did not matter whether the deviation was right or left of the imaginary line.

Horizontal deviation: This metric was calculated analogous to stripe deviation with the difference that the imaginary line was placed horizontal in between the two stripes. Both the directional and absolute horizontal deviation were calculated.

Center deviation: This metric was calculated as the deviation away from the arena center.

All python metrics were calculated in 4 ways: (1) while the animal is moving, (2) while stationary, (3) the combination of 1 and 2 and (4) the last minus an edge correction factor of 11.78% to remove edge artefacts.

#### Repeated behavioral testing

Individual animals were tested repeatedly in the behavioral arena. To do so, animals were after a behavioral test placed back into their individual vial. It was possible to repeatedly test individual animals even over the duration of several weeks. For the prolonged testing, individual flies were flipped every 2-3 days.

#### Correlation between brain properties and behavior

After performing behavioral tests, individual flies were placed back into their individual vial to keep track of their identity. The next day the flies were dissected and fixed (3.7% Formaldehyde in PBS) in individual containers (500μl Eppendorf tubes) for 20 minutes. After several washes with PBS, all individual flies were placed in individual quadrants on poly(L)lysine coated (Sigma, p1524) grid slides (EMS, 63405-01). The identity of the fly in each quadrant was recorded and after performing immunohistochemistry on the slide, each brain hemisphere in every quadrant was imaged using a Nikon TI2000 AIR or Leica SP8 confocal microscope.

After imaging, full reconstructions of neuronal morphology were created semi-automatically using Bitplane Imaris 9.2 and the reconstructions were blindly measured for DCN and Medulla axon numbers. For the number of medulla axons, a line was drawn in the presumptive medulla plate region and all axons were counted that grew from this region into the posterior medulla. After merging the behavioral with the morphological analysis, the data was analyzed using heat maps for correlations between the behavioral and anatomical data. Afterwards, these correlations were further statistically validated using R.

#### Induction of visual asymmetry by single eye blinding

Wildtype flies were raised as in all other experiments and tested in the Buridan arena. After the behavioral test a single fly eye was painted with a lacquer pen with an extra fine tip (Kreul, 0.8 mm black, 47411). After 48h of resting time the same individuals were retested for their single eyed response. The paired data was analyzed using the pairwise Wilcoxon test.

#### Statistics

All data was statistical analyzed using R. First, we investigated normal data distributions using the Shapiro-Wilk test. Then we choose the appropriate parametric or non-parametric test for our data. For normal distributed data we were using an ANOVA followed by the Tukey test for the pairwise comparisons. Data homogeneity was tested using the pairwise F-test. Paired data was analyzed using the paired Wilcoxon test.

For our correlative analyses we were using either the parametric Pearson product-moment correlation coefficient or the non-parametric Spearman’s rank-order correlation coefficient. For the Pearson product-moment correlation we considered an r of 0.0-0.19 as a very weak correlation, from 0.2-0.39 as a weak correlation from 0.4-0.59 as a moderate correlation, from 0.6-0.79 as a strong correlation and from 0.8-1.0 as a very strong correlation(*11*).

---

**Fig. S1.**
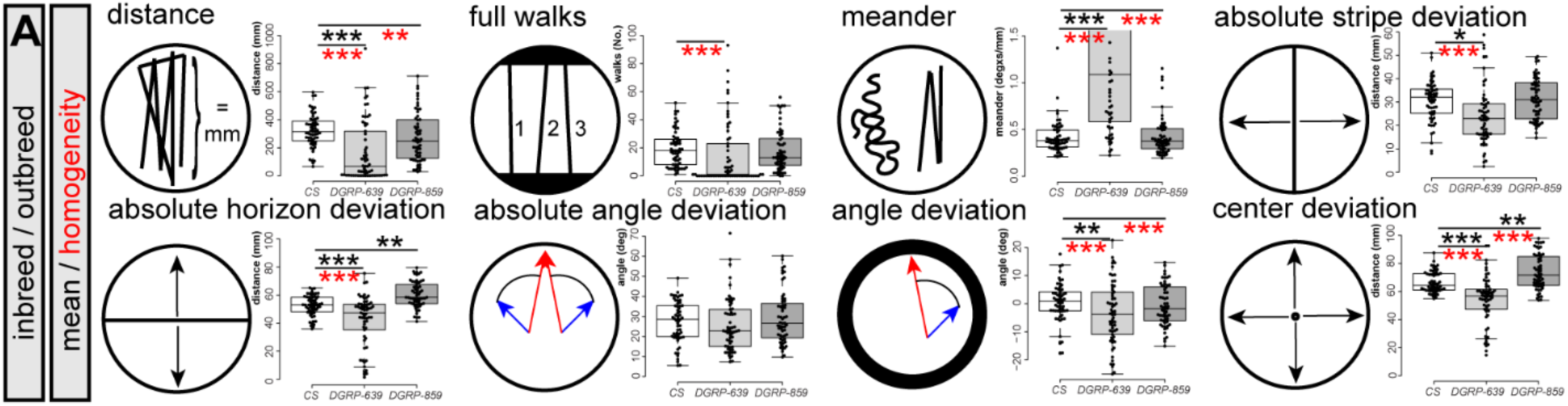
in relation to Fig.1. A) Detailed trajectory analysis reveals that the outbred control (*CS*, N=60) differs on the population level from the inbred lines (*DGRP-639*, N=61, *DGRP-859*, N=59, statistical test: two-way ANOVA and Tukey HSD as post hoc test and pairwise F-test for homogeneity) both in their distributions around the mean and data homogeneity. Nevertheless, despite having less genetic variation, the inbred lines show less homogeneity and they are therefore more variable in most behavioral parameters. *DGRP-639* flies walk significantly shorter distances than *CS* flies (p-value < 0.001). Both DGRP lines show increased variability in the distance compared to *CS* (*CS* – *DGRP-639*, p-value < 0.001; *CS* – *DGRP-859*, p-value = 0.003). The number of full walks between the stripes is comparable for all three genotypes, only the data distribution is different for *CS* and *DGRP-639* (p-value < 0.001). The straightness of the path or meander is different between *CS* and *DGRP-639* (p-value < 0.001). The data distribution differs between *CS* and both DGRP lines (*CS* – *DGRP-639*, p-value < 0.001; *CS* – *DGRP-859*, p-value < 0.001). The mean and homogeneity of absolute stripe deviation differs for *CS* and *DGRP-639* (p-value = 0.01, p-value < 0.001). The absolute horizon deviation differs in the mean for *CS* and both DGRP lines (*CS* – *DGRP-639*, p-value < 0.001; *CS* – *DGRP-859*, p-value = 0.002). DGRP-639 shows a more spread data distribution than CS (p-value < 0.001). The absolute angle deviation is neither significantly different for the data mean or homogeneity. The angle deviation corrected differs significantly for the mean of *CS* and *DGRP-639* (p-value = 0.03). Additionally, the data distribution differs for *CS* and both DGRP lines (*CS* – *DGRP-639*, p-value < 0.001; *CS* – *DGRP-859*, p-value < 0.001). The center deviation significantly differs both in the mean (*CS* – *DGRP-639*, p-value < 0.001; *CS* – *DGRP-859*, p-value = 0.009) and homogeneity (*CS* – *DGRP-639*, p-value<0.001; *CS* – *DGRP-859*, p-value < 0.001) for *CS* and both DGRP lines.

**Fig. S2.**
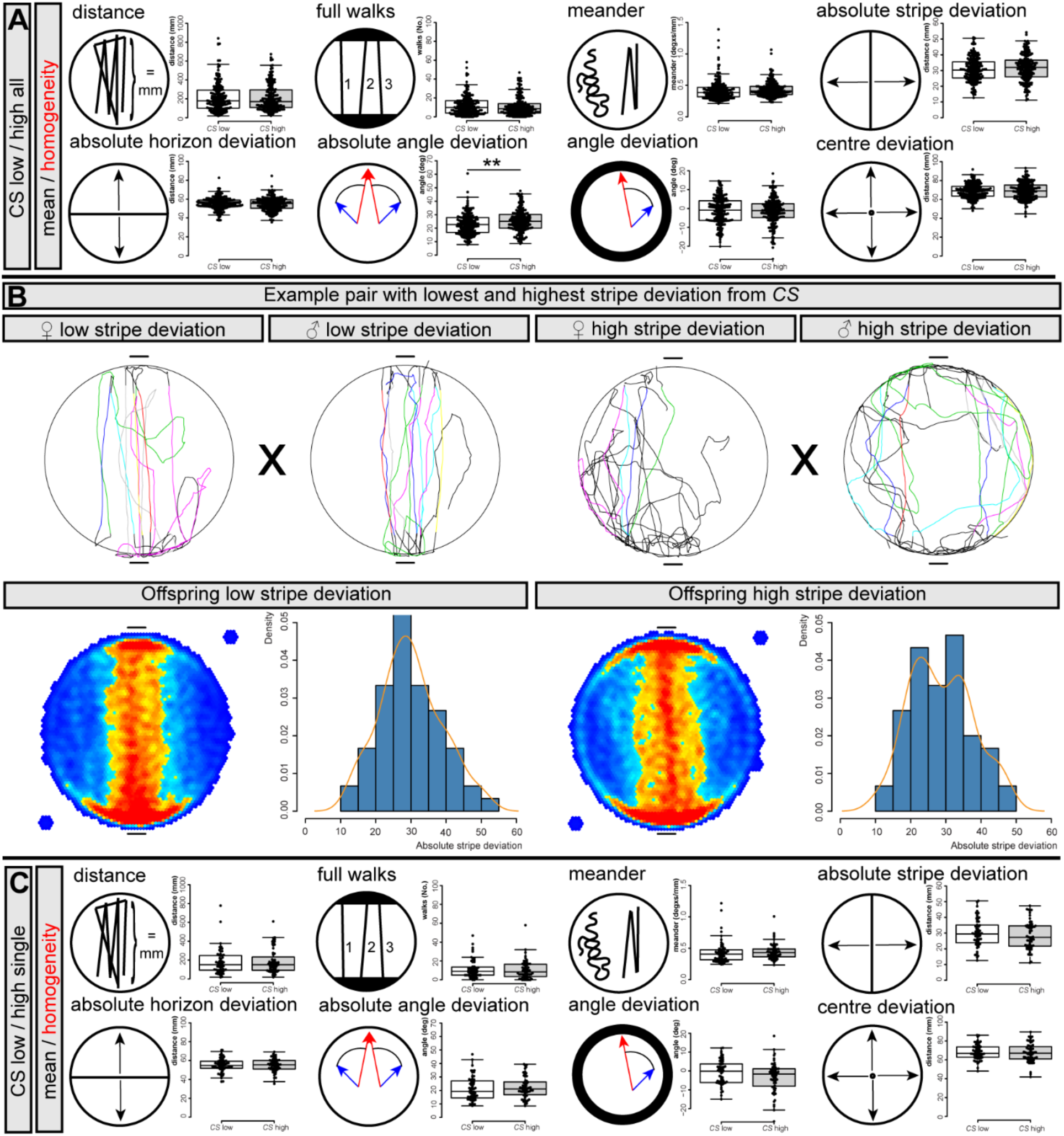
in relation to Fig.2. A) Statistical analysis for the *CS* selection experiment reveals that the offspring is behaving identically (N=180, 180). No differences were found in mean and homogeneity (two-way ANOVA and Tukey HSD as post hoc test and pairwise F-test for homogeneity) for the distance (p-value = 0.9), full walks (p-value = 0.5), meander (p-value = 0.5), absolute stripe deviation (p-value = 0.22), absolute horizon deviation (p-value = 0.5), angle deviation corrected (p-value = 0.7), center deviation (p-value = 0.7). Only the absolute angle deviation was statistically different between the two groups (p-value = 0.002). B) A single best and worst couple out of the selection experiment from 47 *CS* males and 37 virgin *CS* females. The path plots show in the top row from left to right the virgin female with the lowest stripe deviation, the male with the lowest stripe deviation, the virgin female with the highest stripe deviation and the male with the highest stripe deviation. The offspring of these two parental couples is shown in the bottom row. Neither the heatmap nor the histogram display any of two populations (N=60, 60) different from each other. Also, the statistical analysis for stripe deviation shows with a p-value of 0.4 (two-way ANOVA and Tukey HSD as post hoc test) no statistical difference between the two populations. C) Further statistical analysis for the selection experiment in B) reveals that the offspring (N=60, 60, two-way ANOVA and Tukey HSD as post hoc test) is behaving identical in all parameters (distance, p-value = 0.7; full walks, p-value = 0.6; meander, p-value = 0.6; absolute stripe deviation, p-value = 0.4; absolute angle deviation, p-value 0.5; angle deviation corrected, p-value = 0.2; center deviation, p-value = 0.6).

**Fig. S3.**
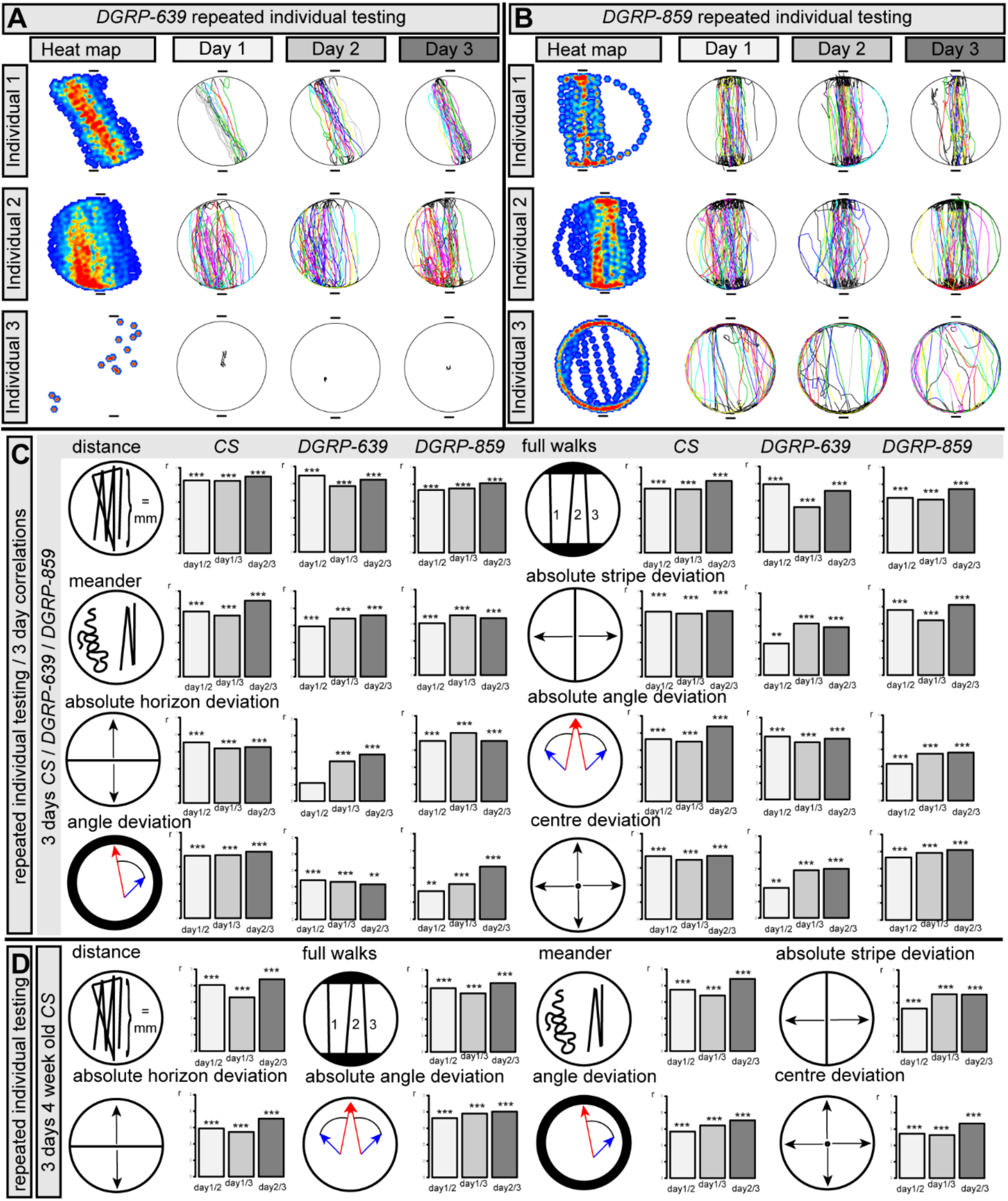
in relation to Fig.3. A) Adult *DGRP-639* animals (N=52) were repeatedly tested over the duration of three days. The flies show remarkably stability in their responses independent of whether they respond strong or weakly to the visual cue or even show no response towards the visual cue at all. The heatmap and the individual tracks for the three days show very persistent behavior. The top row shows an animal with left shifted strong behavior, the middle row an animal with weaker response that predominantly stays at the bottom left half of the arena. The lower row is an example of the in this genotype frequently found flies that hardly move at all. B) Adult *DGRP-859* flies (N=76) were repeatedly tested over the duration of 3 days. The flies show remarkably stability in their responses independent of whether they respond strong or weakly to the visual cue or even show no response towards the visual cue at all. The heatmap and the individual tracks for the three days show very persistent behavior. The top row shows an animal with straight strong behavior, the middle row an animal with a weaker stripe response and a wider path in between the stripes. The lower row is an example for an edge centric fly. C) Statistical analysis for *CS* (N=74), *DGRP-639* (N=52), *DGRP-859* (N=76) shows that the responses for many behavioral parameters are highly correlated (Pearson correlation coefficient). The total distance is highly correlated for all three genotypes: *CS*: Day 1 vs. Day 3: r=0.85, p-value<0.001, Day 1 vs. Day 3: r=0.85, p-value<0.001, Day 2 vs. Day 3: r=0.9, p-value<0.001; *DGRP-639*: Day 1 vs. Day 2: r=0.89, p-value<0.001, Day 1 vs. Day 3: r=0.77, p-value<0.001, Day 2 vs. Day 3: r=0.84, p-value<0.001; *DGRP-859*: Day 1 vs. Day 2: r=0.73, p-value<0.001, Day 1 vs. Day 3: r=0.75, p-value<0.001, Day 2 vs. Day 3: r=0.81, p-value<0.001. The number of full walks in between the stripes is similarly highly correlated: *CS*: Day 1 vs. Day 2: r=0.75, p-value<0.001, Day 1 vs. Day 3: r=0.75, p-value<0.001, Day 2 vs. Day 3: r=0.84, p-value<0.001; *DGRP-639*: Day 1 vs. Day 2: r=0.79, p-value<0.001, Day 1 vs. Day 3: r=0.52, p-value<0.001, Day 2 vs. Day 3: r=0.71, p-value<0.001; *DGRP-859*: Day 1 vs. Day 2: r=0.64, p-value<0.001, Day 1 vs. Day 3: r=0.62, p-value<0.001, Day 2 vs. Day 3: r=0.74, p-value<0.001. The straightness of the path or meander is also highly correlated for all 3 genotypes: *CS*: Day 1 vs. Day 2: r=0.76, p-value<0.001, Day 1 vs. Day 3: r=0.72, p-value<0.001, Day 2 vs. Day 3: r=0.89, p-value<0.001; *DGRP-639*: Day 1 vs. Day 2: r=0.59, p-value<0.001, Day 1 vs. Day 3: r=0.68, p-value<0.001, Day 2 vs. Day 3: r=0.72, p-value<0.001; *DGRP-859*: Day 1 vs. Day 2: r=0.61, p-value<0.001, Day 1 vs. Day 3: r=0.70, p-value<0.001, Day 2 vs. Day 3: r=0.67, p-value<0.001. The absolute stripe deviation is especially highly correlated for *CS* and *DGRP-859*. *DGRP-639* is less strongly correlated for Day 1 vs. Day 2: *CS*: Day 1 vs. Day 2: r=0.76, p-value<0.001, Day 1 vs. Day 3: r=0.74, p-value<0.001, Day 2 vs. Day 3: r=0.72, p-value<0.001; *DGRP-639*: Day 1 vs. Day 2: r=0.39, p-value=0.0047, Day 1 vs. Day 3: r=0.63, p-value<0.001, Day 2 vs. Day 3: r=0.59, p-value<0.001; *DGRP-859*: Day 1 vs. Day 2: r=0.76, p-value<0.001, Day 1 vs. Day 3: r=0.64, p-value<0.001, Day 2 vs. Day 3: r=0.82, p-value<0.001. The absolute horizon deviation is similarly to the absolute stripe less strongly correlated for *DGRP-639* especially for Day 1 vs. Day 2: *CS*: Day 1 vs. Day 2: r=0.71, p-value<0.001, Day 1 vs. Day 3: r=0.64, p-value<0.001, Day 2 vs. Day 3: r=0.66 p-value<0.001; *DGRP-639*: Day 1 vs. Day 2: r=0.22, p-value=0.12,, Day 1 vs. Day 3: r=0.49, p-value<0.001, Day 2 vs. Day 3: r=0.57, p-value<0.001; *DGRP-859*: Day 1 vs. Day 2: r=0.71, p-value<0.001, Day 1 vs. Day 3: r=0.80, p-value<0.001, Day 2 vs. Day 3: r=0.71, p-value<0.001. The absolute angle deviation is highly correlated between days for all genotypes. The Pearson correlation coefficient is lowest for *DGRP-859*: *CS*: Day 1 vs. Day 2: r=0.71, p-value<0.001, Day 1 vs. Day 3: r=0.64, p-value<0.001, Day 2 vs. Day 3: r=0.66 p-value<0.001; *DGRP-639*: Day 1 vs. Day 2: r=0.77, p-value<0.001, Day 1 vs. Day 3: r=0.70 p-value<0.001, Day 2 vs. Day 3: r=0.74, p-value<0.001; *DGRP-859*: Day 1 vs. Day 2: r=0.43, p-value<0.001, Day 1 vs. Day 3: r=0.55, p-value<0.001, Day 2 vs. Day 3: r=0.56, p-value<0.0013. The angle deviation is for all genotypes correlated between the days. *CS* exceeds the correlations of the two DGRP lines: *CS*: Day 1 vs. Day 2: r=0.73, p-value<0.001; Day 1 vs. Day 3: r=0.74, p-value<0.001, Day 2 vs. Day 3: r=0.78 p-value<0.001; *DGRP-639*: Day 1 vs. Day 2: r=0.48, p-value<0.001, Day 1 vs. Day 3: r=0.46, p-value<0.001, Day 2 vs. Day 3: r=0.43, p-value=0.002; *DGRP-859:* Day 1 vs. Day 2: r=0.33, p-value=0.004, Day 1 vs. Day 3: r=0.41, p-value<0.001, Day 2 vs. Day 3: r=0.61, p-value<0.001. All three genotypes show strong correlations between days for center deviation. The lowest correlation is found for *DGRP-639* between Day1 and Day 2: *CS*: Day 1 vs. Day 2: r=0.74, p-value<0.001, Day 1 vs. Day 3: r=0.69, p-value<0.001, Day 2 vs. Day 3: r=0.74 p-value<0.001; *DGRP-639*: Day 1 vs. Day 2: r=0.37, p-value=0.007, Day 1 vs. Day 3: r=0.58, p-value<0.001, Day 2 vs. Day 3: r=0.6, p-value<0.001; *DGRP-859*: Day 1 vs. Day 2: r=0.73, p-value<0.001, Day 1 vs. Day 3: r=0.78, p-value<0.001, Day 2 vs. Day 3: r=0.82, p-value<0.001. D) Statistical analysis for aged four-week old *CS* (N=74) shows that also for aged flies behavioral responses are highly conserved throughout the 3-day period. The correlation results are statistically identical to the results from younger *CS* flies. The total distance walked is highly correlated between the different days: Day 1 vs. Day 2: r=0.81, p-value<0.001, Day 1 vs. Day 3: r=0.66, p-value<0.001, Day 2 vs. Day 3: r=0.88, p-value<0.001. Also, for the number of full walks the results between days are remarkably similar between individual flies: Day 1 vs. Day 2: r=0.78, p-value<0.001, Day 1 vs. Day 3: r=0.72, p-value<0.001, Day 2 vs. Day 3: r=0.84, p-value<0.001. The same is true for the meander of the path of individual flies: Day 1 vs. Day 2: r=0.75, p-value<0.001, Day 1 vs. Day 3: r=0.68, p-value<0.001, Day 2 vs. Day 3: r=0.88, p-value<0.001. The responses for absolute stripe deviation are highly correlated: Day 1 vs. Day 2: r=0.53, p-value<0.001, Day 1 vs. Day 3: r=0.7, p-value<0.001, Day 2 vs. Day 3: r=0.7, p-value<0.001. Also, the responses for the absolute horizon deviation are significantly correlated: Day 1 vs. Day 2: r=0.59, p-value<0.001, Day 1 vs. Day 3: r=0.55, p-value<0.001, Day 2 vs. Day 3: r=0.71 p-value<0.001. Absolute angle deviations are also highly correlated for aged individuals between days: Day 1 vs. Day 2: r=0.72, p-value<0.001, Day 1 vs. Day 3: r=0.78, p-value<0.001, Day 2 vs. Day 3: r=0.80 p-value<0.001. Similar is true for angle deviation, with lower correlation coefficients: Day 1 vs. Day 2: r=0.57, p-value<0.001, Day 1 vs. Day 3: r=0.64, p-value<0.001, Day 2 vs. Day 3: r=0.71 p-value<0.001 Also, the center deviation is correlated between days with lower correlation coefficients than found for other parameters: Day 1 vs. Day 2: r=0.54 p-value<0.001, Day 1 vs. Day 3: r=0.53, p-value<0.001, Day 2 vs. Day 3: r=0.67 p-value<0.001

**Fig. S4.**
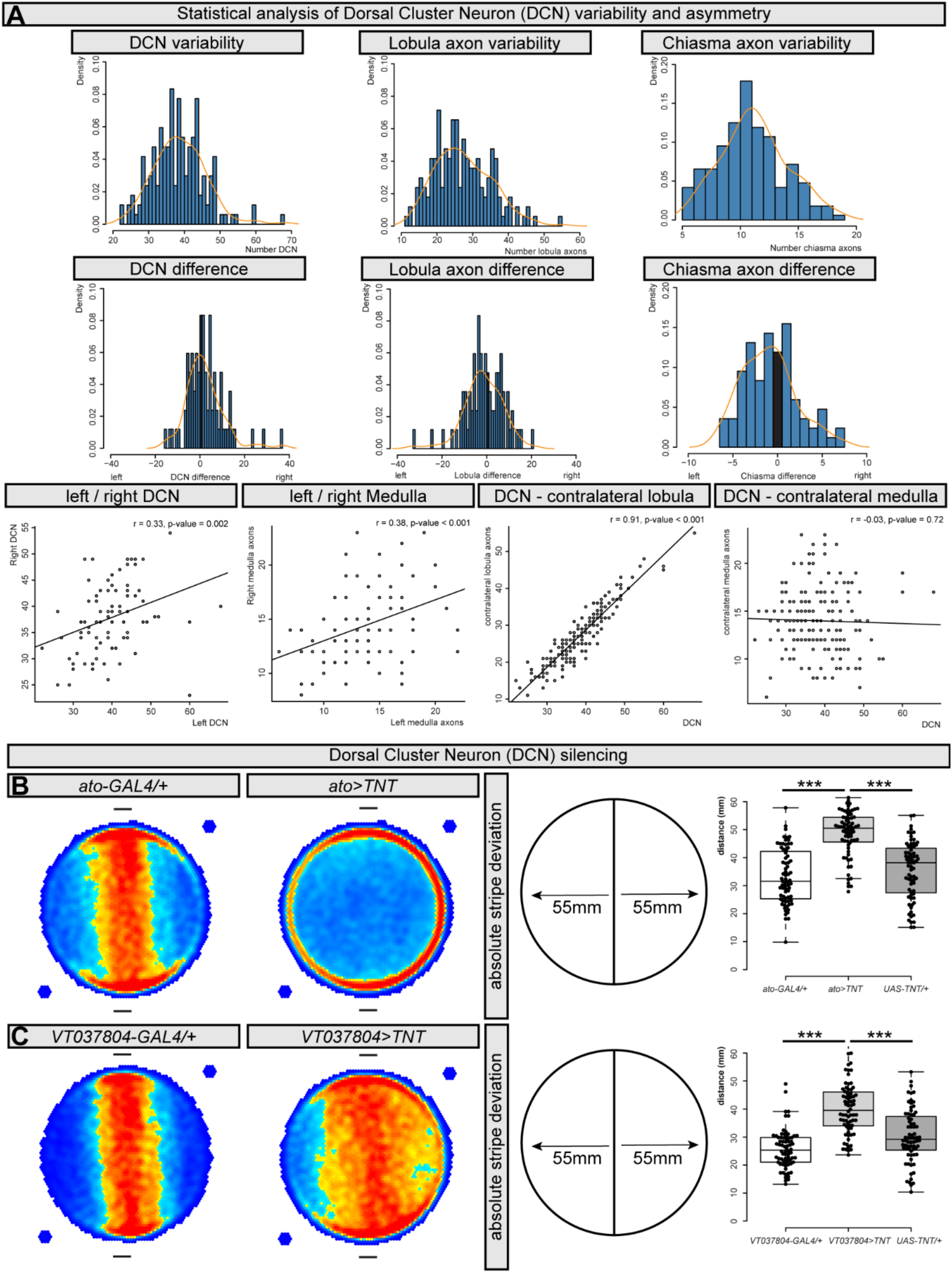
in relation to Fig.4. A) Statistical analysis of DCN variability (N = 168), asymmetry (N = 84) and right-left correlativity (N = 84). The number of DCNs varies as shown in the histogram between 22 to 68 cell bodies per brain hemisphere. The mean number of DCNs is 38.74. The number of axons in the lobula varies from 11 to 55 axons with a mean of 27.48. The number of axons exiting the lobula / lobula plate range from 5 to 19 axons as measured in the chiasma with a mean of 11.27. The DCN asymmetry between left and right ranges from 0 to 37 cell bodies with a mean of 5.96. The lobula asymmetry ranges from 0 to 33 axons with a mean of 6.67. The medulla axon asymmetry measured in the chiasma ranges from 0 to 8 with a mean of 2.44. The right and left DCN cell numbers correlate with a Pearson correlation coefficient of r = 0.33 with a p-value = 0.002. The right and left medulla correlate with an r = 0.38 and a p-value < 0.001. The number of DCN cell bodies very highly correlates with the number of lobula axons (r = 0.91, p-value < 0.001. The number of DCN cell bodies does not correlate with the number medulla axons (r = −0.03, p-value = 0.72). B) DCN neuron silencing leads to a complete loss of the stripe fixation response. The heatmap of the control population of *ato-GAL4/+* flies show a normal response in the two-stripe arena. This is entirely lost upon silencing of DCN neurons in *ato>TNT* animals. Statistical analysis (N = 71 Set) of the absolute stripe deviation shows that the *ato>TNT* animals show significant higher stripe deviation than the controls (p-value < 0.001). Higher stripe deviation means that the animals fixate the stripes less. C) Similar results are obtained by silencing DCN neuron silencing with *VT037804-GAL4.* The heatmap of the control population of *VT037804-GAL4/+* flies show a normal response in the two-stripe arena. This is lost upon silencing of DCN neurons in *VT037804>TNT* animals. Statistical analysis (N=71 Set) of the absolute stripe deviation shows that the *VT037804>TNT* animals show significant higher stripe deviation than the controls (p-value < 0.001).

**Fig. S5.**
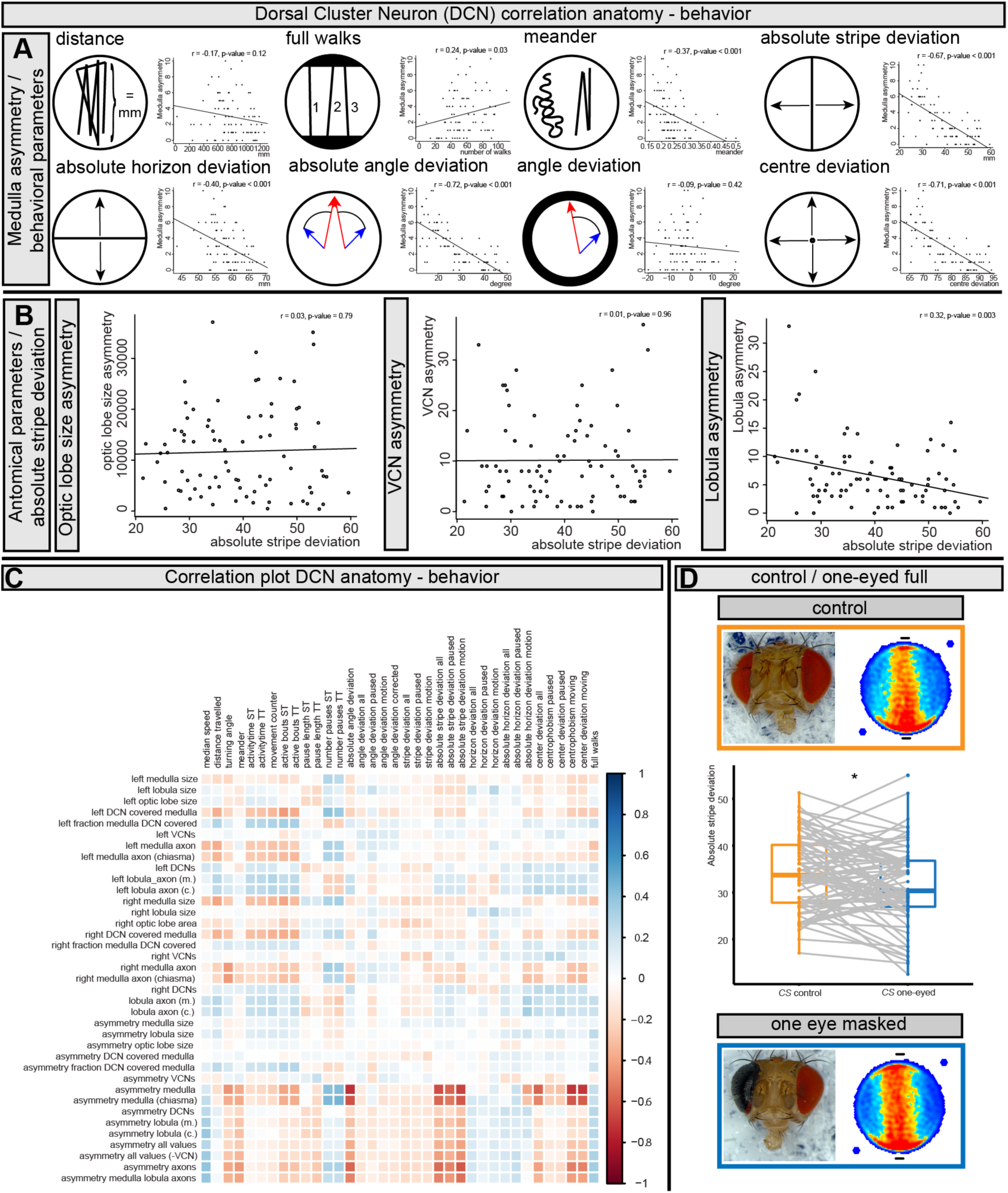
in relation to Fig.5. A) Comparison of the medulla asymmetry and eight representative behavioral parameters (N = 103, Pearson correlation coefficients). The distance the flies walk very weakly negatively correlated with the medulla asymmetry (r = −0.17, p-value = 0.12). The number of full walks between the stripes weakly correlated with the medulla asymmetry (r= 0.24, p-value = 0.03). Similarly, the meander of the tracks weakly negatively correlated with the medulla asymmetry (r = −0.37, p-value <0.001). The absolute stripe deviation strongly negatively correlated with medulla asymmetry (r = −0.67, p-value < 0.001). The absolute horizon deviation moderately negatively correlated with medulla asymmetry (r = −0.40, p-value < 0.001). The absolute angle deviation strongly negatively correlated with medulla asymmetry (r = 0.72, p-value < 0.001). The angle deviation very weakly negatively correlated with medulla asymmetry (r = −0.09, p-value = 0.42). The center deviation strongly negatively correlated with medulla asymmetry (r = 0.71, p-value < 0.001). B) Comparison of a selection of anatomical parameters with absolute stripe deviation (N = 103). The optic lobe size asymmetry very weakly correlated with the absolute stripe deviation (r = 0.03, p-value = 0.79). Also the asymmetry of the other population of *atonal* positive cells (VCN) only correlated very weakly with the absolute stripe deviation (r = 0.01, p-value = 0.96). The Lobula axon asymmetry weakly correlated with absolute stripe deviation (r = 0.32, p-value = 0.003) C) The correlation plot shows a selection of relevant comparisons between DCN anatomy and behavior using the Pearson correlation coefficient (values range from 1 = very strongly correlated to −1 = very weakly correlated, N = 103). D) Full dataset for the paired experiment where flies were repeatedly tested after blinding them (N = 79). Single eyed-blinding resulted in a mild reduction of absolute stripe deviation (p-value = 0.01, paired Wilcoxon test).

**Fig. S6.**
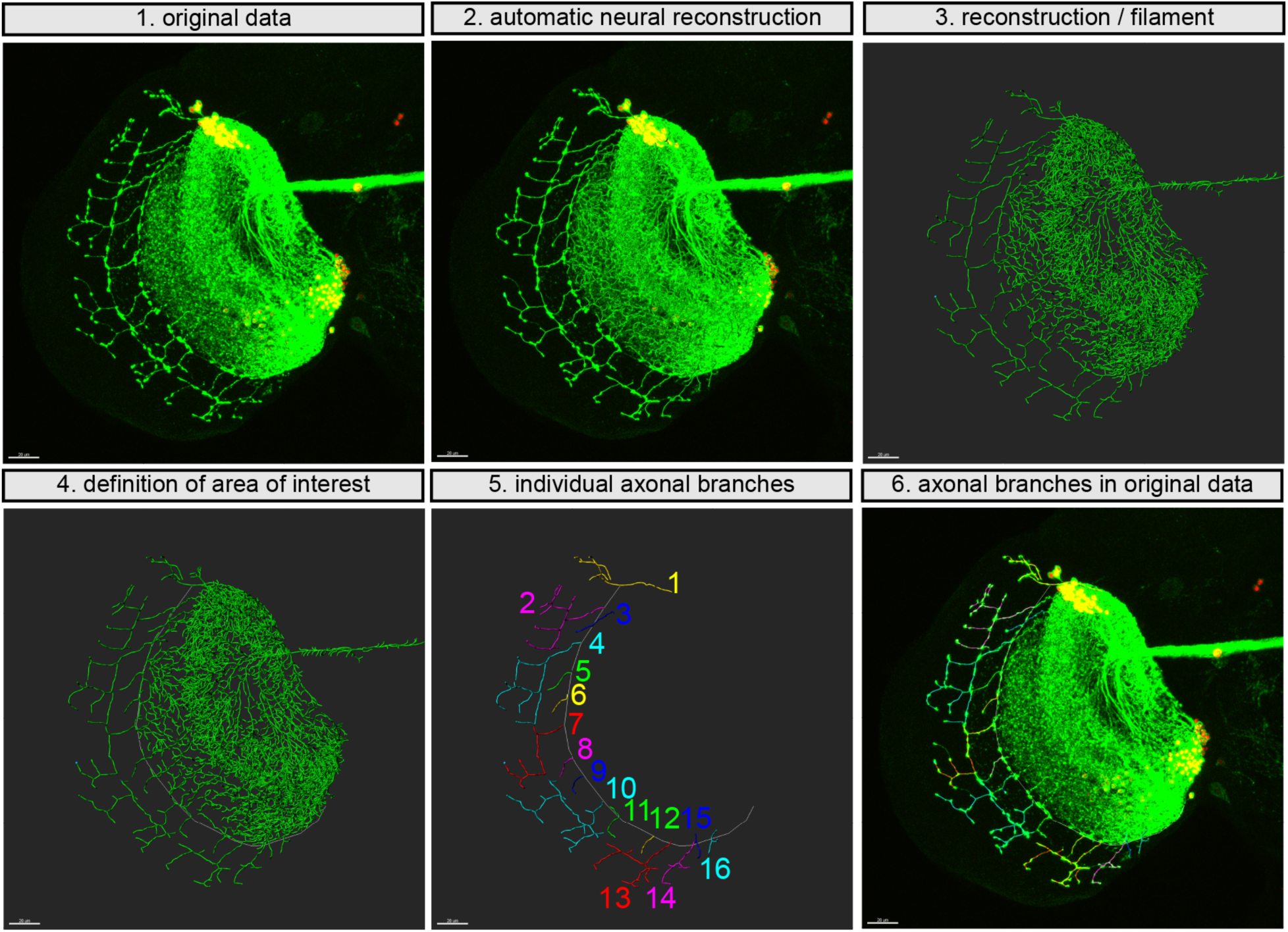
in relation to Fig.5. Workflow of the anatomical reconstruction of medulla axons illustrated for a single side. 1. The original data is imported into Bitplane Imaris 9.2. 2.-3. The neuronal data is reconstructed using the filament function to reconstruct all neural branches. 4. The area of interest in this case the medulla branches is defined by the line separating the lobula plate from the medulla via an optical chiasma (shown in white). 5. Individual axonal branches in the area of interest are reconstructed from the filament. 6. Individual filaments are shown with the original data in the background. Scale bars correspond to 20μm.

**Movie S1 in relation to Fig. 1**

A wildtype *CS* fly with a low stripe deviation score. The 5 min movie shows the platform marked in yellow and the fly tracks in red.

**Movie S2 in relation to Fig. 1**

A wildtype *CS* fly with an intermediate stripe deviation score. The 5 min movie shows the platform marked in yellow and the fly tracks in red.

**Movie S3 in relation to Fig. 1**

A wildtype *CS* fly with a high stripe deviation score. The 5 min movie shows the platform marked in yellow and the fly tracks in red.

**Movie S4 in relation to Fig. 4**

The movie shows the location of the DCNs in the anterior visual field that corresponds to the most posterior medulla and anterior lobula. The GFP marked DCNs are shown in green and the neuropiles are marked in red (α-N-Cadherin).

**Movie S5 in relation to Fig. 4**

The movie shows the location of the DCNs in the anterior visual field on the left side of one individual. The DCNs are marked in green and the postsynaptic cells to the DCNs are marked in red using Trans-Tango(*49*).

**Movie S6 in relation to Fig. 4**

The movie shows the location of the DCNs in the anterior visual field on the right side of one individual. The DCNs are marked in green and the postsynaptic cells to the DCNs are marked in red using Trans-Tango(*49*).

**Movie S7 in relation to Fig. 4**

The movie shows that the DCN medulla branches as marked with GFP (in grey values) in the adult show small movements but overall their pattern is highly stable and does not change over several hours (as shown in this movie) and days.

